# The *las* and *rhl* Quorum Sensing Systems in *Pseudomonas aeruginosa* Form a Multi-Signal Reciprocal Network Which Can Tune Reactivity to Variations in Physical and Social Environments

**DOI:** 10.1101/2023.02.23.529764

**Authors:** Stephen Thomas, Ayatollah Samir El-Zayat, James Gurney, Jennifer Rattray, Sam P. Brown

## Abstract

Researchers often view the multi-signal quorum sensing systems of *Pseudomonas aeruginosa* as a hierarchy, topped by the *las* system which acts as a master regulator. By experimentally controlling the concentration of auto-inducer signals in a signal null strain (PAO1*ΔlasIΔrhlI*), we show that the two primary quorum sensing systems—*las* and *rhl*—act reciprocally rather than hierarchically. Just as the *las* system’s 3-oxo-C_12_-HSL can induce increased expression of *rhlI*, the *rhl* system’s C_4_-HSL increases the expression level of *lasI*. We develop a mathematical model to quantify relationships both within and between the *las* and *rhl* quorum sensing systems and the downstream genes they influence. The results show that not only do the systems interact reciprocally, but they do so cooperatively and nonlinearly, with the combination of C_4_-HSL and 3-oxo-C_12_-HSL increasing expression level far more than the sum of their individual effects. We computationally assess how our parameterized model responds to variation in social (population density) and physical (mass transfer) environment and demonstrate that a reciprocal architecture is more responsive to density and more robust to mass transfer than a strict hierarchy.

## Introduction

Bacterial cells of many species communicate with each other by exchanging diffusible signal molecules. This mechanism, known as quorum sensing (QS), has been well-studied at the level of specific molecular interactions. We now understand how those interactions shape the creation of and response to signal molecules in model organisms such as *Pseudomonas aeruginosa*. We have identified downstream effector genes such as virulence factors whose production depends on QS signals, and we have recognized that some species possess multiple QS circuits. Despite this knowledge, we face gaps in our understanding of how quorum sensing influences bacterial behavior. How does QS guide bacterial actions in response to environmental conditions? What benefits do multiple QS circuits provide? And ultimately, how does QS contribute to bacterial fitness? Answering these questions requires an understanding of quorum sensing at the dynamical systems level as well as the molecular level.

Quorum sensing relies on several components interacting in a dynamical system. Individual cells synthesize small molecules called signals or inducers. These diffuse or are actively transported between the intracellular and extracellular environments. Within cells, signal molecules bind to receptor proteins forming transcription factors. As signal concentration grows, genes activated by these transcription factors trigger a change in the cell’s behavior. Those components related to a particular signal molecule—the signal synthase, the signal molecule, and the cognate receptor—form a quorum sensing system. Some bacterial species have multiple QS systems, the opportunistic pathogen *Pseudomonas aeruginosa* among them. Its *las* and *rhl* acyl-homoserine lactone (AHL) signaling systems have been especially well studied. The *las* system includes the LasI synthase, N-(3-oxododecanoyl)-l-homoserine lactone (3-oxo-C_12_-HSL) signal, and LasR receptor. The corresponding components of the *rhl* system are RhlI, N-butyryl-homoserine lactone (C_4_-HSL), and RhlR. Schuster and Greenberg (2007) estimate that these two systems control expression of as much as 10% of the bacterial genome.

*P. aeruginosa* provides a model for understanding interactions between multiple QS systems. Can the behavior of one system, determined by the concentration of signal it produces, affect the behavior of a different system, specifically by increasing or decreasing expression of the second system’s synthase or receptor? We classify possible multi-system architectures into three broad patterns shown in Figure 1. *Independent systems* (Figure 1A) have no influence on each other; *hierarchical systems* (Figure 1B) have a relationship but only in one direction, and *reciprocal systems* (Figure 1C) each exert influence on the other. At this level we do not consider the underlying mechanism(s) of the inter-system effects. For example, the signal of one system may bind directly to the receptor of the other; alternatively, the signal/receptor complex of one system may act as a transcriptional regulator of components in the second system. In both cases we simply denote the first system as influencing the second.

**Figure 1.**
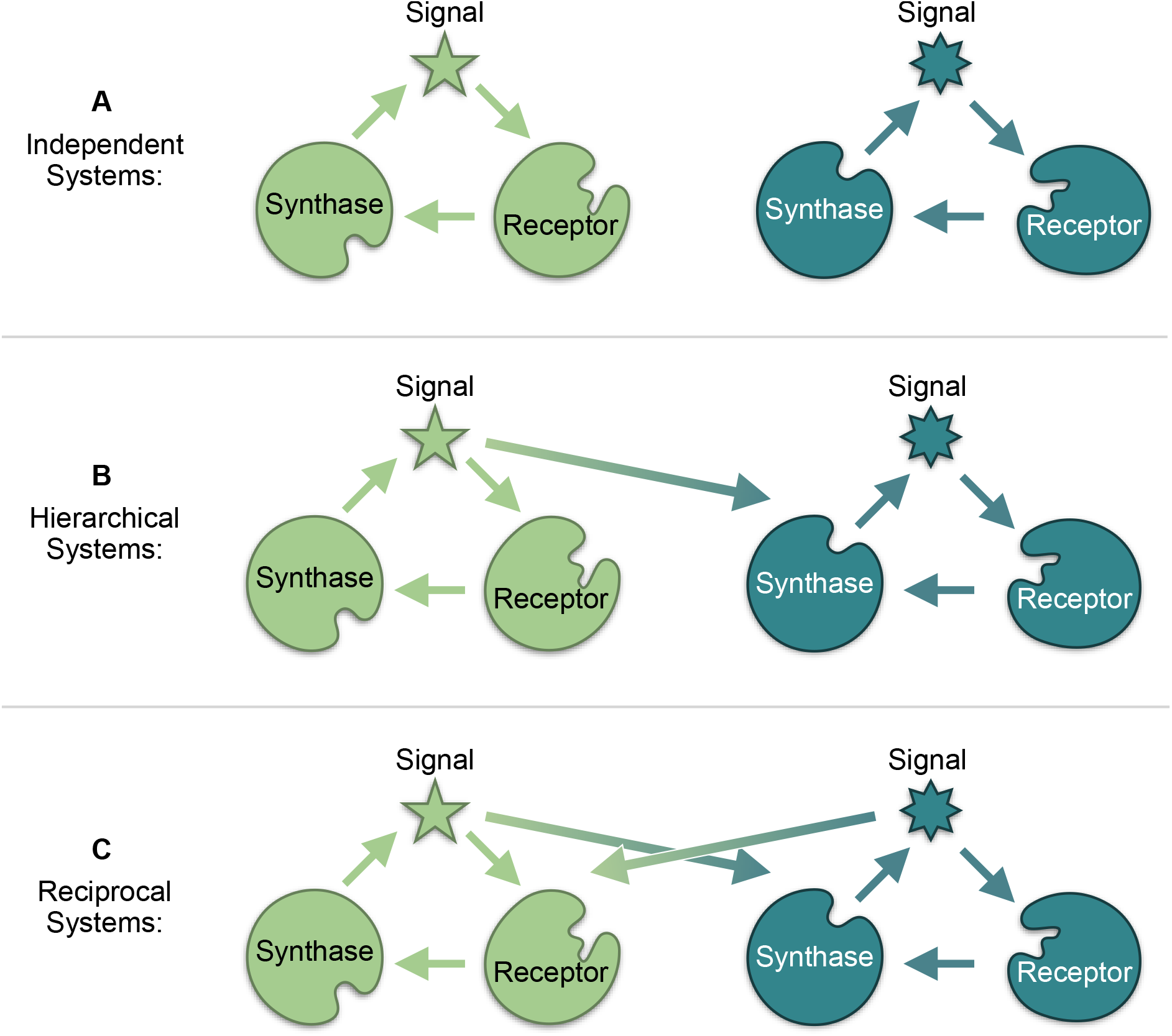
The relationship between quorum sensing systems may be classified as independent, hierarchical, or reciprocal. The top row shows two independent systems: the signal of each has no influence on the expression of synthase or receptor in the other. In the middle row, one system’s signal does influence expression of the other’s components; this system is considered the top of a hierarchical relationship. When both systems’ signals influence the others’ components, as in the bottom row, a reciprocal relationship exists.

In the case of *las* and *rhl*, independent, isolated operation was eliminated as early as 1996 when Latifi et al. used *lacZ* transcriptional fusions to show that the combination of LasR and 3-oxo-C_12_-HSL controls expression of *rhlR*, demonstrating that the *las* system influences the *rhl* system. These and other results have led many researchers to view *las* and *rhl* as a hierarchy, with the *las* system serving as master QS system controlling both its own activation and that of the *rhl* system (Figure 2). We confirm this consensus view via a structured literature review (Tables SI.1 and SI.2). The review literature is silent on whether the *rhl* signal C_4_-HSL, either alone or when bound to RhlR, can influence the expression of the *las* synthase or receptor. There are however, reports of exactly this behavior in the primary literature (Wellington and Greenberg 2019; Jayakumar et al. 2022). If *lasI* or *lasR* also respond to RhlR, then the strict hierarchical view may be missing an important factor that determines the overall system response.

**Figure 2.**
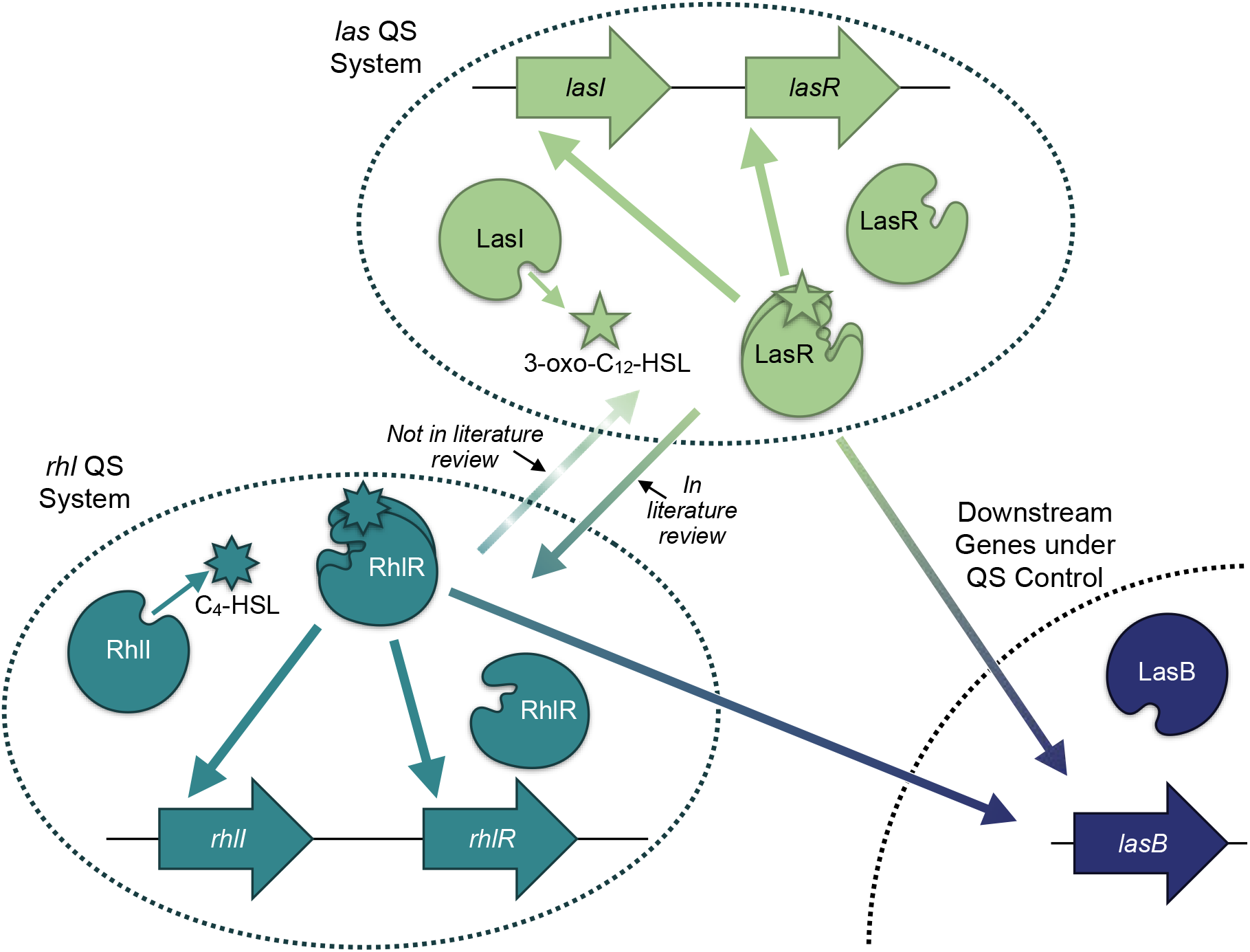
The *P. aeruginosa* QS regulatory network is typically viewed as a hierarchy, with the *las* system on top. Solid arrows summarize the relationships depicted in 17 review papers published since 1996 (Tables SI.1 and SI.2). All papers show the *las* system affecting the *rhl* system, but none identify a *las* synthase or receptor gene as a target of the *rhl* system (dashed line).

Our experiments seek to address this omission, and the resulting data reveal three key insights. First, the traditional hierarchical view of *las* and *rhl* is incomplete. Our results confirm that *las* can exert control over the *rhl* system. In addition, however, we observe the converse: *rhl* influences the *las* system, specifically expression of *lasI*. Second, we show that maximum expression of genes in both QS systems requires both signals in combination. Finally, we demonstrate that this architecture can make QS-controlled behavior more sensitive to population density and more robust to interfering physical environmental conditions.

## Results

To uncover interactions between the *las* and *rhl* systems, we experimentally assess QS gene expression in a signal null strain (PAO1*ΔlasIΔrhlI*) exposed to defined, exogenous concentrations of the signal molecules C_4_-HSL and 3-oxo-C_12_-HSL. We use bioluminescence (lux) reporters for *lasI* and *rhlI* to estimate expression levels of the respective genes. We then develop mathematical models to quantify the effects of each system on the other and their consequent responses to environmental variation.

### The *las* and *rhl* Systems Influence Each Other

We first evaluate quorum sensing behavior under the influence of a single signal. We establish a baseline expression level by measuring reporter luminescence with no signal present. We then observe the increase in luminescence as exogenously controlled signal concentration increases. The ratio of luminescence with signal to luminescence with no signal represents the fold-change in expression induced by the defined signal concentration. Figure 3 shows the results for 3-oxo-C_12_-HSL. As expected, expression of both genes increases as signal concentration increases. The availability of the *las* signal molecule influences the expression of *rhlI* as well as *lasI*, and, therefore, the *las* system affects the *rhl* system.

**Figure 3.**
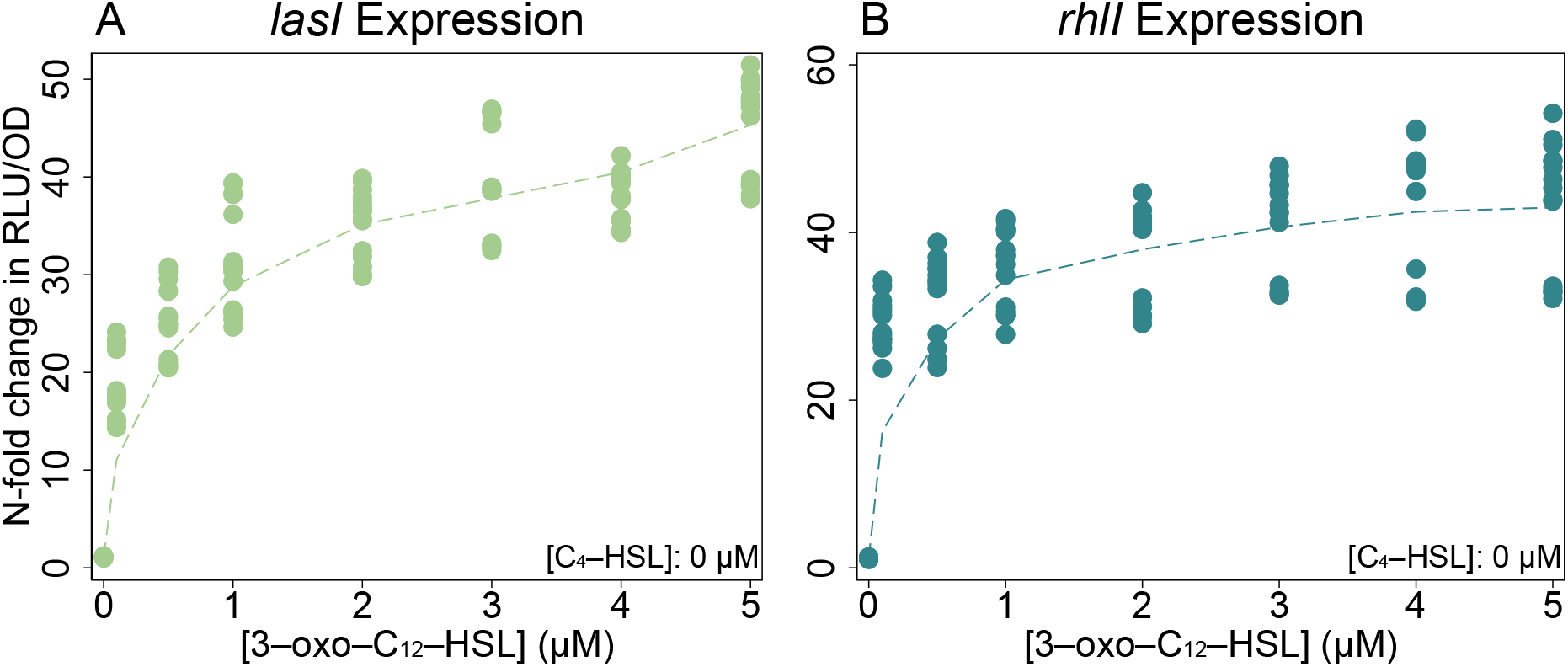
The *las* signal 3-oxo-C_12_-HSL increases the expression of *lasI* and *rhlI* in a signal null PAO1. Plots show fold-change in RLU/OD (relative light units per optical density) values compared to baseline with no exogenous signals in NPAO1*ΔlasIΔrhlI* cultures. Genomic reporter fusions *lasI:luxCDABE, rhlI:luxCDABE*, and *lasB:luxCDABE* were used to generate luminescence. Points are individual observations within the time window of peak expression; dashed lines show a locally weighted regression of the mean fold-change for each concentration value.

While we find no surprises with 3-oxo-C_12_-HSL, our experiments with C_4_-HSL challenge the conventional hierarchical view. Figure 4 shows those results: expression of *las* and *rhlI* increases with higher C_4_-HSL concentration. The response of *lasI* (Figure 4A) does not correspond to a simple hierarchy with *las* as the master. Here we find that the *rhl* system affects the *las* system.

**Figure 4.**
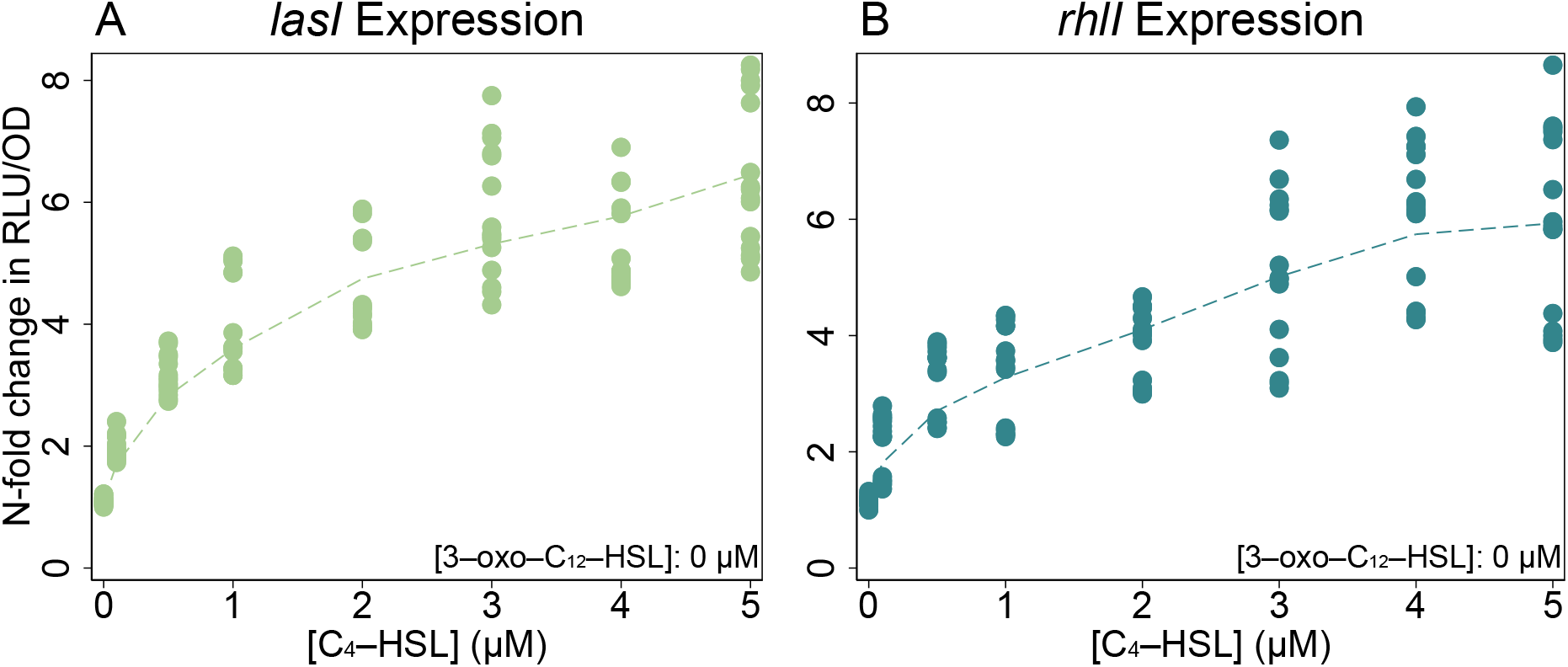
The *rhl* signal C_4_-HSL increases the expression of *lasI* and *rhlI* in a signal null PAO1. Expression of both genes grows with increased concentration of C_4_-HSL. Plots are constructed as in Figure 3. Strains and reporters also as in Figure 3. Note that panel A shows results are not captured by the consensus *las→rhl* hierarchy, as it clearly indicates that *lasI*, in the *las* system, responds to the signal produced by the *rhl* system.

To quantify the impact of each signal alone, we model gene expression using Michaelis-Menten kinetics under quasi-steady state assumptions. The resulting dynamics provide a simple model of transcription factor binding (Santillán 2008; Boluri 2008 chapter 9) as well as more general processes such as enzyme activity and substrate-receptor binding.

For a single signal, gene expression is defined by equation 1, where *a0* is basal expression, *a* is the maximum increase in expression from auto-induction, [*S*] is the signal concentration, and *K* is the disassociation constant of the binding event or, equivalently, the signal concentration corresponding to half of the maximum expression gain. With this model we quantify two qualities: how strongly a signal can increase gene expression above its basal level (*a*), and how sensitive gene expression is to the presence of the signal (*K*).

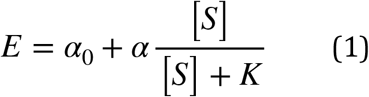

By minimizing the sum of squared error, we estimate model parameters from our data, using only those observations in which a single signal is present. Table 1 presents the results as maximum fold-change and half-concentration values for both signals. Our model fits illustrate that while the *las* and *rhl* systems have reciprocal impacts, those impacts are not symmetrical. The *las* signal 3-oxo-C_12_-HSL has a substantially greater influence on gene expression than C_4_-HSL. In both cases the potential fold-change from 3-oxo-C_12_-HSL is approximately six times greater than the potential fold-change from C_4_-HSL. Both *lasI* and *rhlI* are also more sensitive to 3-oxo-C_12_-HSL than to the C_4_-HSL (by factors of 4 and 30, respectively).

**Table 1.** Single Signal Parameter Estimates. Estimated fold-change, derived from raw parameters of equation 1 as (*a* + *a*_*0*_) / *a*_*0*_, and half-concentration, *K*, values for gene expression as a function of a single signal in isolation. Values shown with 95% confidence intervals.

| Signal | Parameter | *lasI* Estimate | *rhlI* Estimate |
| --- | --- | --- | --- |
| C<sub>4</sub>-HSL | Max fold-change | 6.4 × (5.8 - 7.0) | 6.4 × (5.3 - 7.4) |
|  | ½ conc. | 1.0 µM (0.7 - 1.4) | 1.6 µM (0.8 - 2.4) |
| 3-oxo-C<sub>12</sub>-HSL | Max fold-change | 38 × (36 - 40) | 35 × (34 - 36) |
|  | ½ conc. | 0.24 µM (0.17 - 0.30) | 0.052 µM (0.031 - 0.073) |

### *las* and *rhl* Combine Synergistically

Figures 3 and 4 consider the effects of each signal in isolation, but wildtype cells with functioning synthase genes can produce both signals. To understand environments where both signals are present, we use controlled concentrations of both signals in combination. Figure 5 presents those results in the form of heat maps. The qualitative responses of both genes are similar: raising the concentration of either signal increases expression regardless of the concentration of the other signal. As with our observations of C_4_-HSL alone, these results, in particular the behavior of *lasI*, demonstrate again that the *rhl* system affects the *las* system.

**Figure 5.**
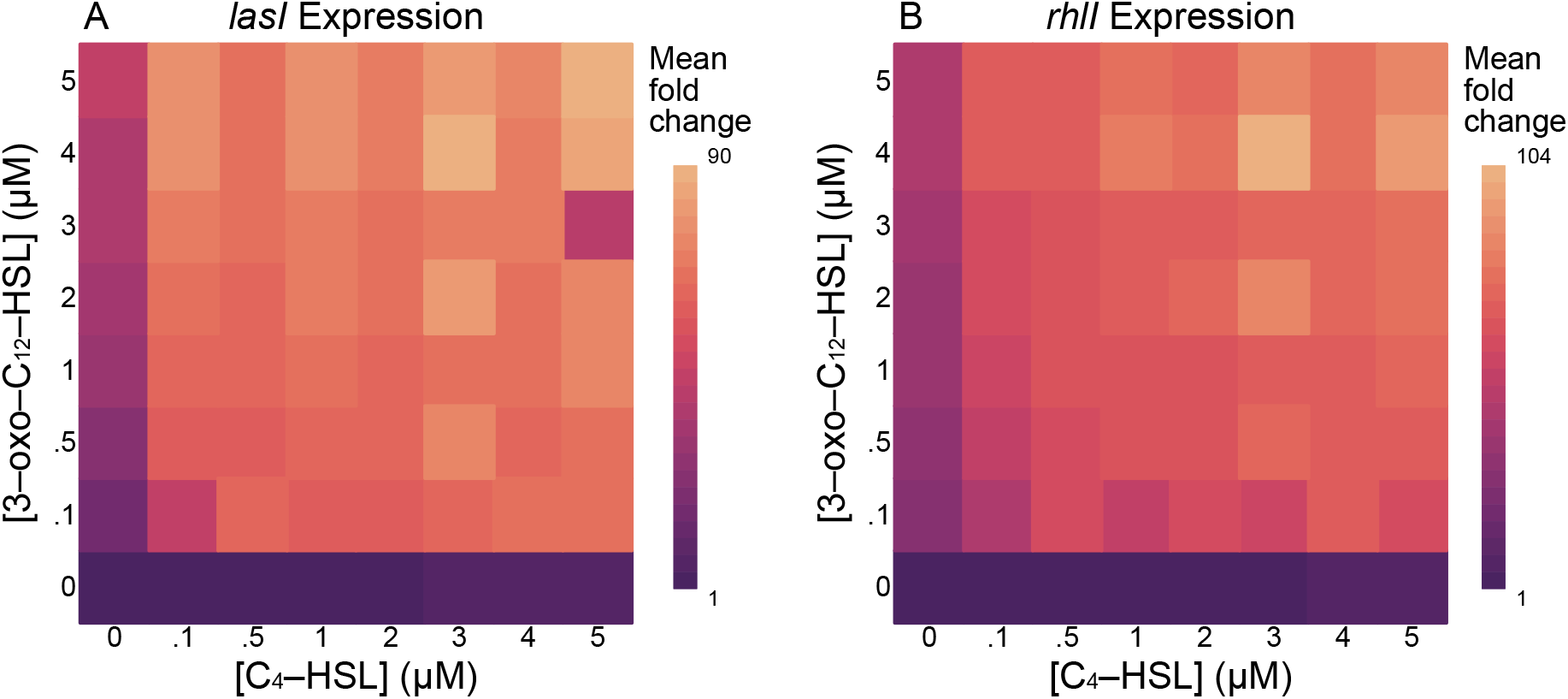
Expression of *lasI* and *rhlI* is maximal in the presence of both C_4_-HSL and 3-oxo-C_12_-HSL. Expression of both genes grows with increased concentration of either signal when both signals are combined. Heatmaps show fold-change in RLU/OD values compared to baseline with no exogenous signals.

Having established a simple model for each signal in isolation, we next consider whether that model is sufficient to explain the effect of the signals in combination. Can we estimate total expression as the sum of expression induced by each signal alone? Such a response could result from two independent binding sites in the promoter regions (Buchler et al. 2003), one site for LasR/3-oxo-C_12_-HSL and a separate site for RhlR/C_4_-HSL. Figure 6 clearly shows that we cannot. The maximum expression observed, shown as a “ceiling” in that figure’s panels, far exceeds the sum of the signals’ individual influence. The presence of both signals boosts expression by as much as 30-fold beyond the level of what a simple sum would predict.

**Figure 6.**
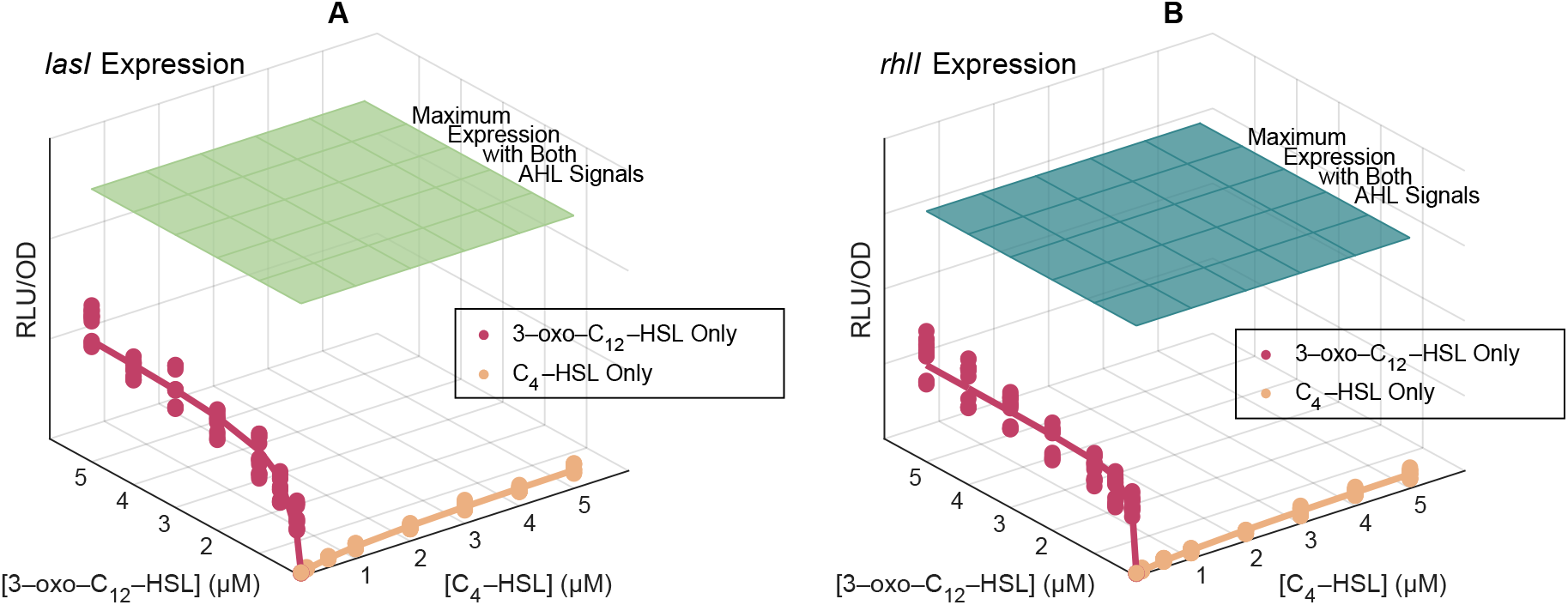
Neither 3-oxo-C_12_-HSL nor C_4_-HSL alone can effect maximal expression of *lasI* or *rhlI*. Both genes require non-zero concentrations of both signals to achieve maximum expression. The flat surfaces in the plots indicate the maximum mean expression level measured across all combinations of signal concentrations. The plotted points represent observed expression levels when C_4_-HSL is withheld (red) and when 3-oxo-C_12_-HSL is withheld (yellow). Lines indicate the model predictions (Equation 1, parameters in Table 1).

To account for the synergy between the signals, we incorporate a cooperativity term in the model. Note that the cooperativity term is a multiplication of signals, and it alone cannot explain the full response, as the product is necessarily zero when any signal is absent. This term accounts for any non-adaptive interaction, for example the ability of one bound transcription factor to recruit the binding of a second transcription factor (Kaplan et al. 2008). Equation 2 shows the result. Each of the three genes has a basal expression level, amplification from each signal alone, and additional amplification from each pair-wise combination of signals. The interaction from these pair-wise combinations captures the cooperative enhancement from the combined signals.

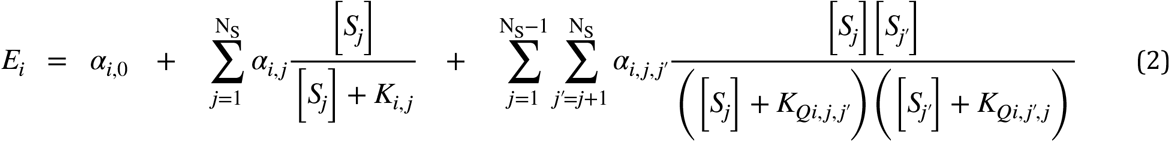

For both sets of observations we again minimize the sum of squared errors to estimate parameters for this multi-signal model. The results of Table 2 show the influence of cooperativity on expression levels. For *lasI* and *rhlI* the maximum expression induced by both signals nearly doubles compared to the maximum expression induced by 3-oxo-C_12_-HSL alone.

**Table 2.** Multi-signal parameter estimates. Model parameters for gene expression as a function of multiple signal concentrations. Parameters are the same as in Table 1 with addition of cooperative fold-change, again derived from raw parameters as (*ai*_,*j,j’*_ + *a*_*0*_) / *a*_*0*_, and cooperative half-concentration *K*_*Q,i,j,j’*_. Values shown with 95% confidence intervals.

| Signal | Parameter | *lasI* Estimate | *rhlI* Estimate |
| --- | --- | --- | --- |
| C<sub>4</sub>-HSL | Max fold-change | 6.4 × (5.8 - 7.0) | 6.4 × (5.3 - 7.4) |
|  | ½ conc. | 1.0 μM (0.7 - 1.4) | 1.6 μM (0.8 - 2.4) |
| 3-oxo-C<sub>12</sub>-HSL | Max fold-change | 38 × (36 - 40) | 35 × (34 - 36) |
|  | ½ conc. | 0.24 μM (0.17 - 0.30) | 0.052 μM (0.031 - 0.073) |
| Combined | Max fold-change | 30 × (29 - 31) | 27 × (26 - 28) |
|  | KQ for C<sub>4</sub>-HSL | 0.003 μM (0 - 0.011) | < 0.001 μM |
|  | KQ for 3-oxo-C<sub>12</sub>-HSL | < 0.001 μM | < 0.001 μM |

Figure 7 summarizes these results graphically. It answers the question posed in Figure 2—the *rhl* system does influence the *las* system—and it shows the relative magnitudes of the effects.

**Figure 7.**
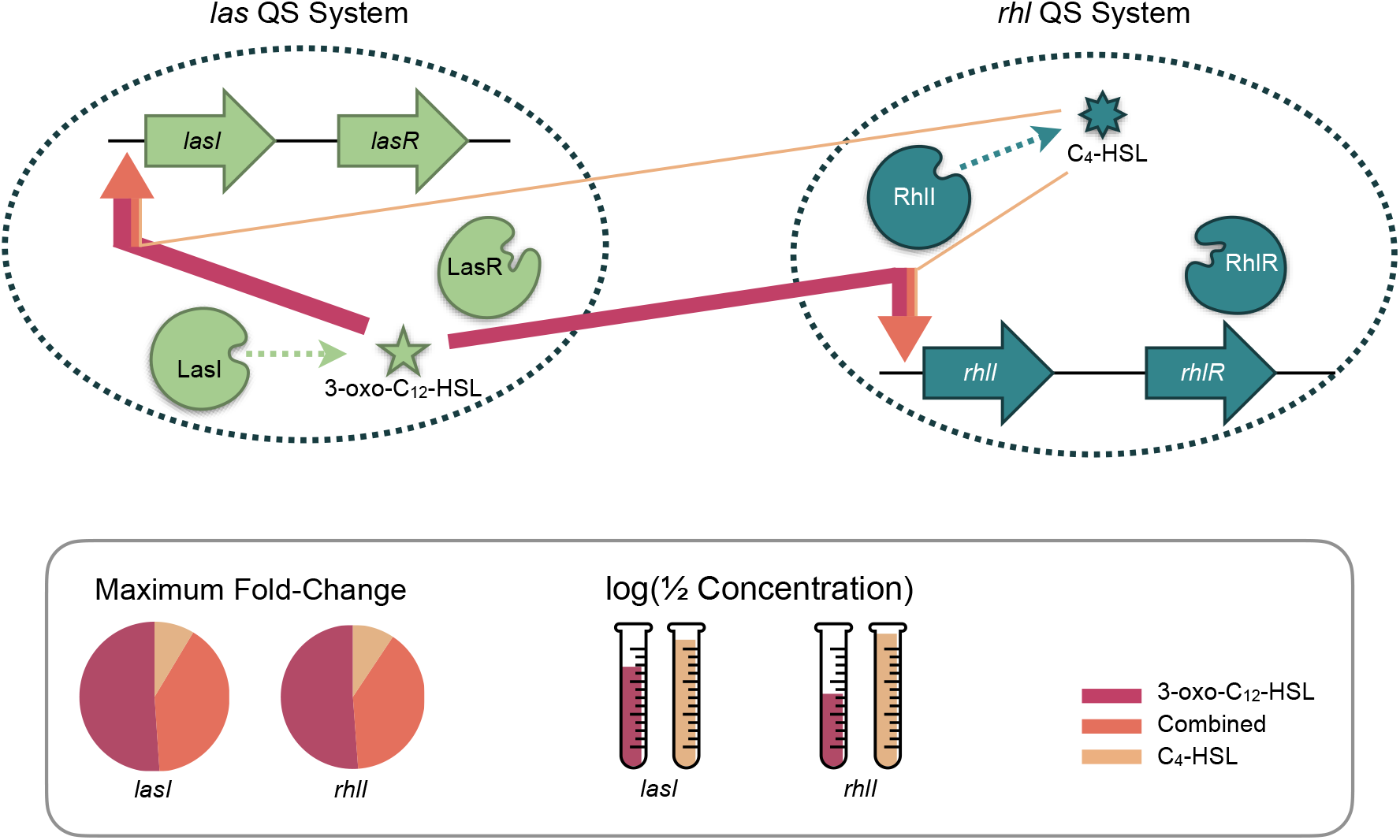
The *las* and *rhl* quorum sensing systems have a reciprocal, but unequal relationship. Red arrows represent maximum fold-change induction from *las*’s 3-oxo-C_12_-HSL and yellow arrows maximum fold-change induction from *rhl’*s C_4_-HSL. The orange component is additional induction from the combination of both signals. Arrow thickness is proportional to fold-change. Inset shows relative contribution of each signal to total maximum fold-change for expression levels of *lasI* and *rhlI*, and half concentration values for each.

### Reciprocity Contributes Significantly to the Quorum Sensing Response

Having established that both signals influence the expression levels of both synthase genes, we next consider how that relationship affects the overall quorum sensing response. That response may be characterized as the extracellular signal concentrations resulting from environmental conditions such as population density. Building on previous models of extracellular signal dynamics (James et al. 2000; Dockery and Keener 2001; Ward et al. 2001; Brown 2013; Cornforth et al. 2014) we assume that signal concentration (a) increases in proportion to the corresponding synthase’s expression level, multiplied by the number of cells expressing synthase, and (b) decreases due to a constant rate of decay. Those assumptions lead to the differential equation model of equation 3, where *S*_*i*_ is the concentration of signal *i, Ei* (**S**) is the expression level of the synthase for signal *i* (as a function of both signal concentrations, **S**) and *c* the proportionality constant, *N* is the population density, and δ*i* is the decay rate of signal *i*.

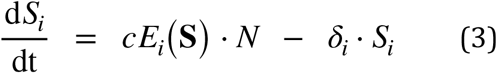

To find solutions for the steady state signal concentrations *Si*\* in this model, we estimate expression levels *Ei* (**S**) from our experimental data (Equation 2, Table 2), approximate the proportionality constant *c* based on data collected for Rattray et al. 2022 (details in supplementary information), and note that the measurements of Cornforth at el. (2014) show the decay rate of 3-oxo-C_12_-HSL (*i* = 1) to be 1.7 times greater than C_4_-HSL (*i* = 2) across a range of environmental conditions. The resulting solutions define the steady state concentrations of both signals as functions of population density for the reciprocal relationship that our data exhibit.

We then compare those results to values that would result from other, hypothetical architectures by adjusting the estimates of synthase expression levels. We do that by setting appropriate interaction coefficients in equation 2 to zero, as detailed in Table 3. Those adjustments allow us to simulate independent and hierarchical architectures using data collected from a reciprocal architecture.

**Table 3.**
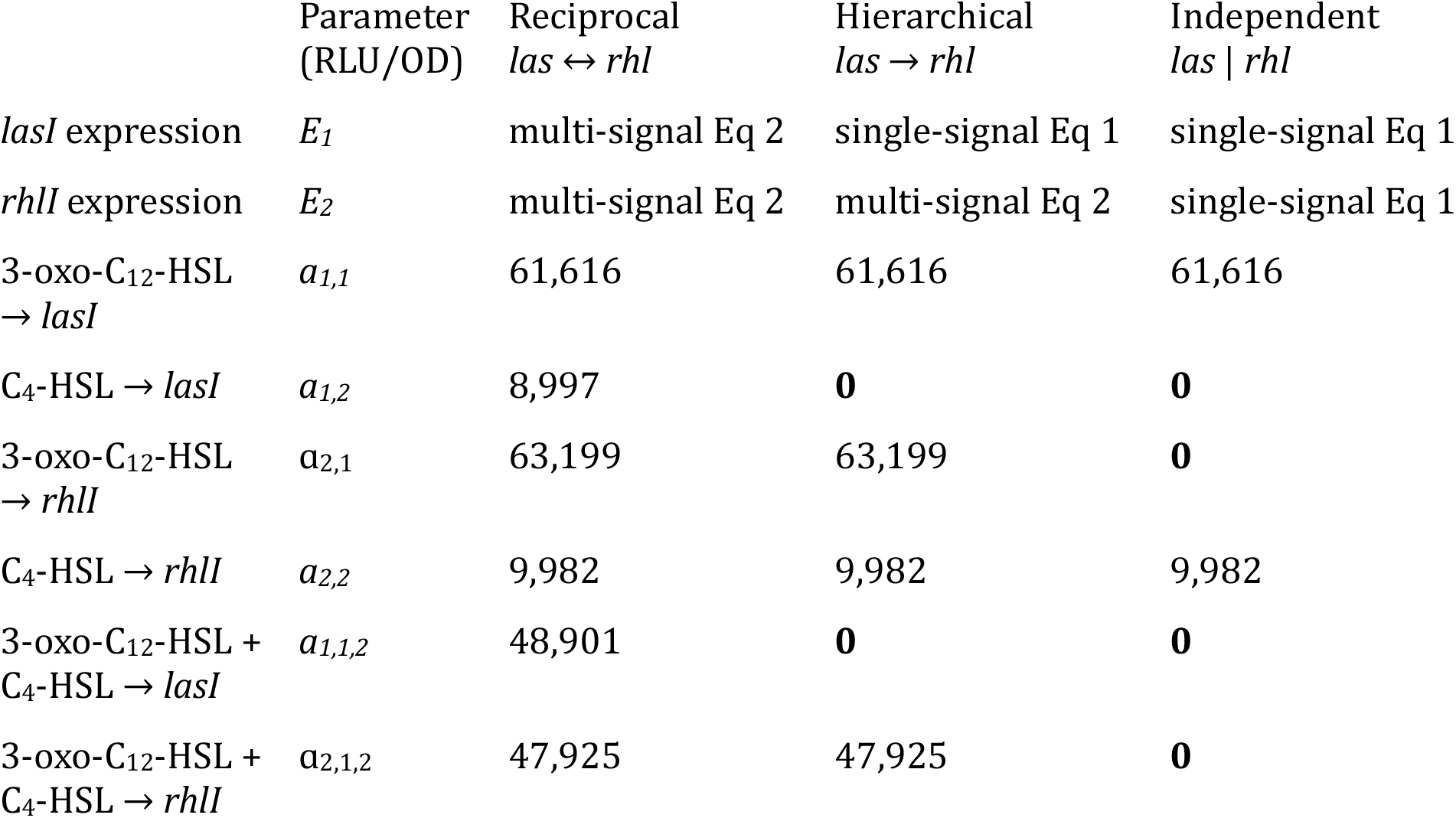
Hierarchical and independent architectures are special cases of the reciprocal architecture with appropriate parameters set to zero. Note that table shows raw parameters from equation 2, in particular, *a* values in units of RLU/OD rather than fold-change (as in Table 2). The values for the reciprocal architecture, however, are equivalent to the fold-change values in Table 2. The hierarchical architecture parameters “zero out” the effect of C_4_-HSL on *lasI*, and the independent architecture parameters eliminate all multi-signal effects.

With these adjustments we can compare the steady state concentrations for reciprocal and hierarchical architectures. Figure 8 shows the results. As expected, the response of C_4_-HSL to density differs little in both architectures, as in both cases the influence of 3-oxo-C_12_-HSL on *rhlI* expression is the same. The 3-oxo-C_12_-HSL response, in contrast, varies significantly. Accounting for the effect of C_4_-HSL on *lasI* expression (the reciprocal architecture) increases the reactivity of 3-oxo-C_12_-HSL to increasing density.

**Figure 8.**
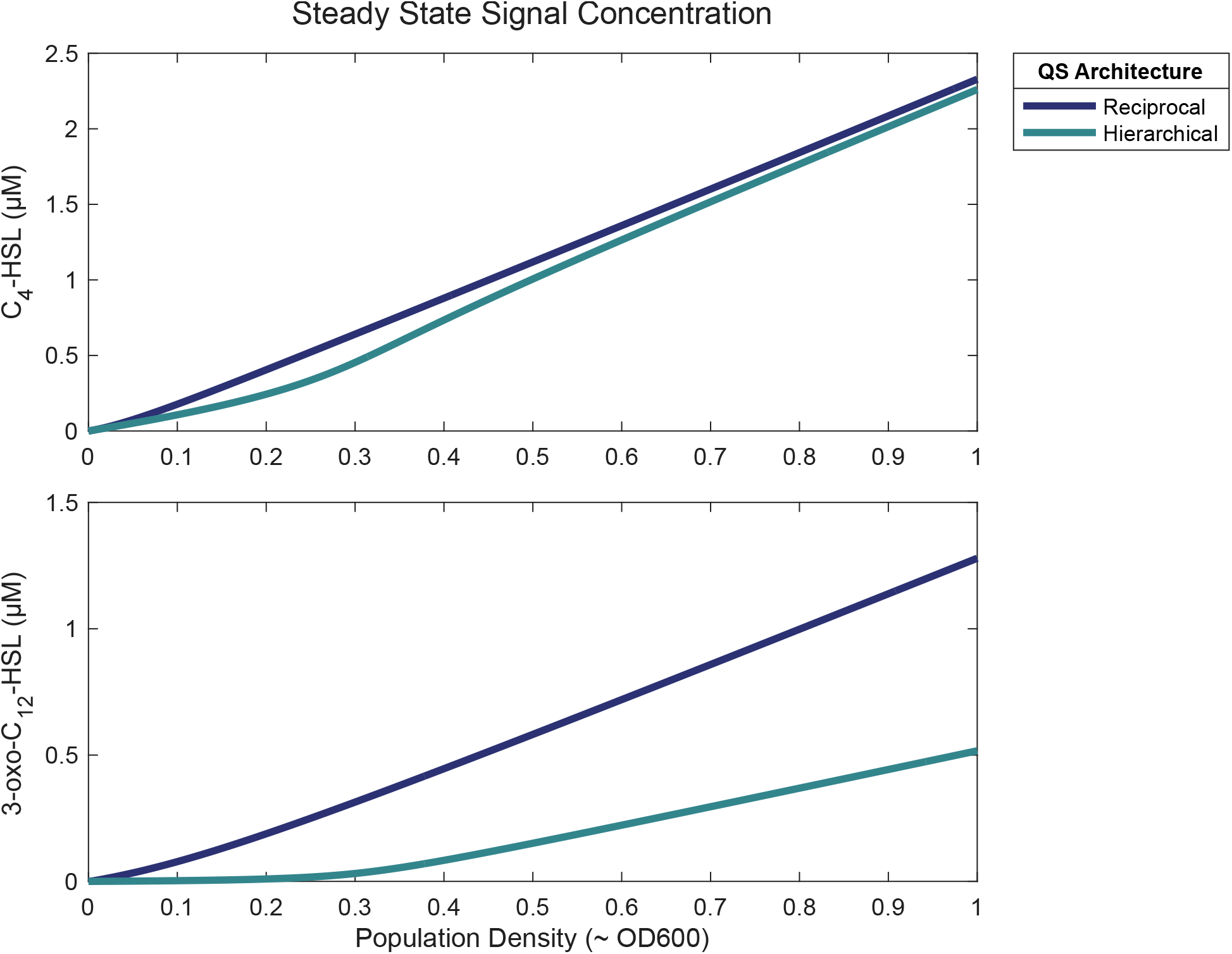
Quorum sensing response varies based on architecture. Plots show the steady concentrations of C_4_-HSL and 3-oxo-C_12_-HSL as a function of population density for both reciprocal and hierarchical architectures. Although C4-HSL values are nearly the same in both architectures, 3-oxo-C12-HSL values differ significantly. Steady state values calculated as equilibrium solutions for equation 3 with parameters from Table 3.

### The Resulting Quorum Sensing Response Shapes Population Behavior

As our interest is ultimately in bacterial behavior in response to quorum sensing, we next consider the expression level of a representative effector gene under quorum sensing control. The *lasB* gene codes for the elastase LasB and is widely used as a model of *P. aeruginosa* virulence (Casilag et al. 2016; Cigana et al. 2021) and cooperation (Diggle et al. 2007; Sexton and Schuster 2017). Significantly, *lasB* expression is known to be influenced by both 3-oxo-C_12_-HSL and C_4_-HSL (Pearson et al. 1997; Nouwens et al. 2003). To quantify that influence we use the same approach as with *lasI* and *rhlI:* measure luminescence of a *lasB* reporter in a signal null strain exposed to defined, exogenous concentrations of both signals. Figure 9 shows the resulting measurements.

**Figure 9.**
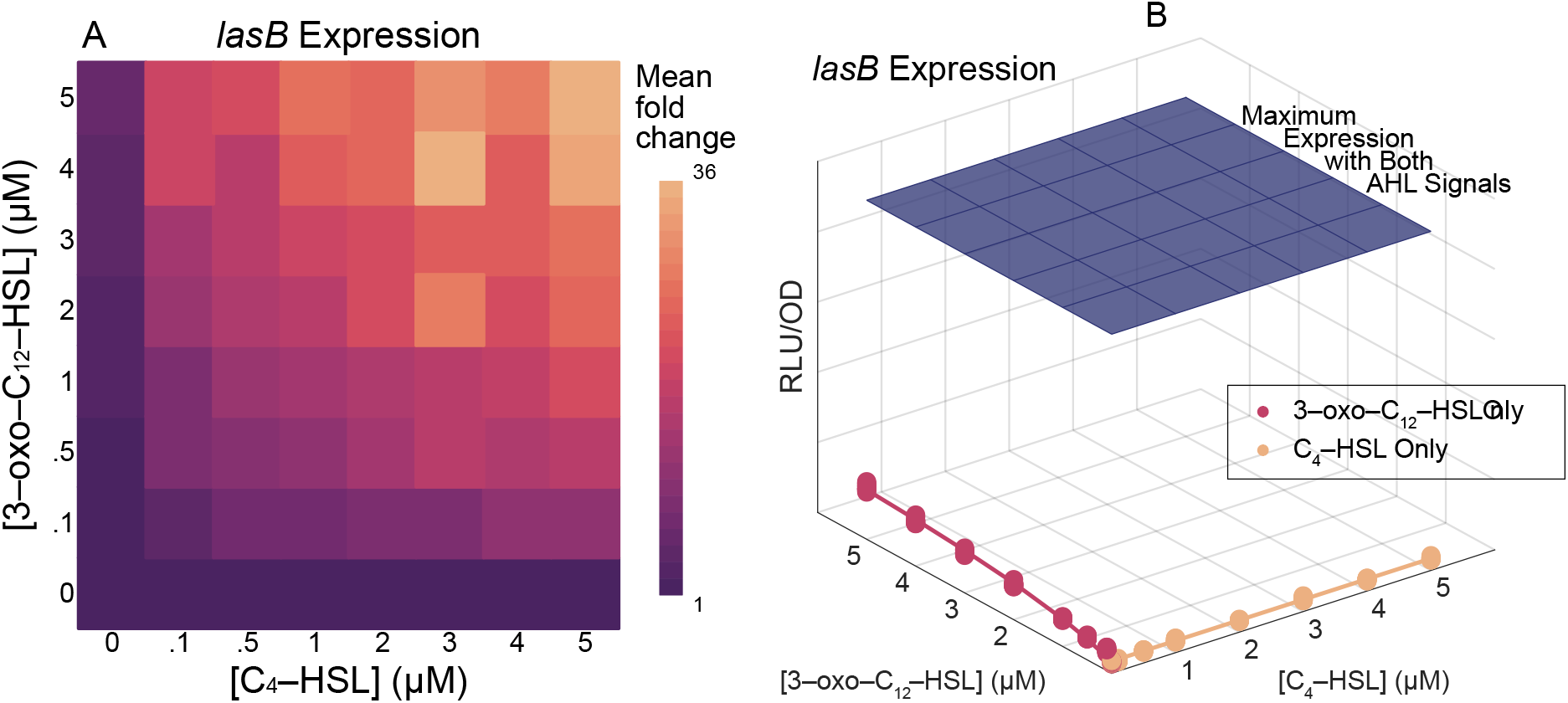
Expression of *lasB* is maximal in the presence of both C_4_-HSL and 3-oxo-C_12_-HSL. Heatmap (panel A) show fold-change in RLU/OD values compared to baseline with no exogenous signals. Surface plot (panel B) shows raw RLU/OD values and compares maximum measured expression (blue “ceiling”) with observed expression levels when C_4_-HSL is withheld (red) and when 3-oxo-C_12_-HSL is withheld (yellow). Lines indicate the model predictions (Equation 1, parameters in Table 4).

These measurements allow us to estimate parameters for a model based on equation 2; Table 4 lists the results.

**Table 4.** Multi-signal parameter estimates for *lasB*. Model parameters for *lasB* expression as a function of multiple signal concentrations. Parameters are the same as in Table 2. Values shown with 95% confidence intervals. Half-concentration estimates less than 0.001 μM are below the limits of precision of the experimental data.

| Signal | Parameter | *lasB* Estimate |
| --- | --- | --- |
| C<sub>4</sub>-HSL | Max fold-change | 1.1 × (1.1 – 1.1) |
|  | ½ conc. | < 0.001 μM |
| 3-oxo-C<sub>12</sub>-HSL | Max fold-change | 6.1 × (5.6 – 6.7) |
|  | ½ conc. | 2.5 μM (1.0 – 3.0) |
| Combined | Max fold-change | 23 × (22 – 24) |
|  | KQ for C<sub>4</sub>-HSL | 0.22 μM (0.18 – 0.25) |
|  | KQ for 3-oxo-C<sub>12</sub>-HSL | 0.42 μM (0.35 – 0.48) |

With the parameter values from table 4 we can predict *lasB* expression for any combination of signal concentrations. In particular, we can use the equilibrium concentrations across the range of environmental conditions as in the example Figure 8. Here we include an additional environmental parameter: advective flow characterized as mass transfer, *m*. The dynamical system, defined in equation 4, can then show variation in response to both social (population density) and physical (mass transfer) environmental variation.

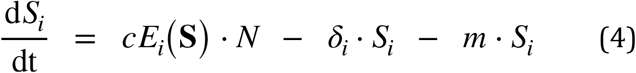

The resulting predictions reveal how those environmental conditions affect *lasB* expression. By repeating this analysis for each of the possible quorum sensing architectures and comparing the results, we show the influence of architecture on QS-controlled behavior.

Figure 10 plots *lasB* expression as reaction norms (Stearns 1989; Rattray et al. 2022) against population density (A, B) and mass transfer rate (C, D) for the three different architectures. As expected, there is a quantitative difference in the behavior of the architectures: in the reciprocal architecture C_4_-HSL increases the expression level of *lasI* and without its influence the *lasB* fold-change is reduced. That overall reduction is evident in the left column (panels A and C). There are also, however, a qualitative differences in the architectures. To show those differences, the right column (panels B and D) artificially scales the hierarchical and independent architectures so that the maximum *lasB* expression matches that of the reciprocal architecture.

**Figure 10.**
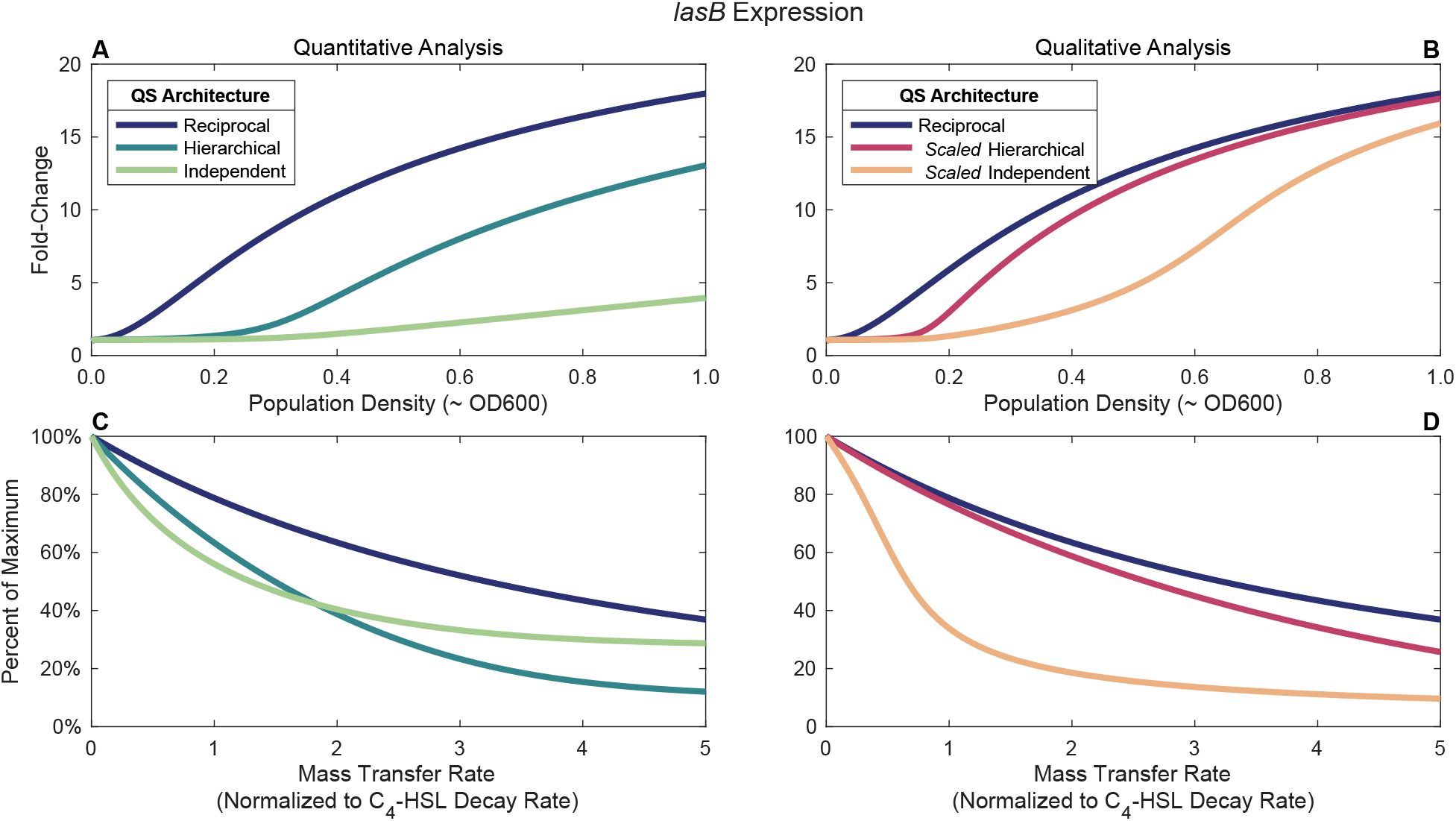
The reciprocal QS architecture is more sensitive to population density and more robust to environmental interference. The graphs show expression of a effector gene under quorum sensing control (*lasB*). All graphs include the results for three QS architectures: independent *las* and *rhl* systems, a hierarchy with the *las* system controlling the *rhl* system, and a reciprocal architecture in which both systems influence each other. Top panels (A and B) show estimated fold-change in expression as a function of population density approximately equivalent to OD600. The range of OD600 values is consistent with Rattray et al. (2022), data from which was used to calibrate parameters. Bottom panels (C and D) show percentage decline from maximum expression as a function of mass transfer rate normalized to C_4_-HSL decay rate. Left panels (A and C) calculated from equation 2 model with parameters from Table 3. Right panels (B and D) emphasize qualitative rather than quantitative differences by scaling the parameters of Table 3 to effect the same maximum *lasI* and *rhlI* expression in all architectures.

The native reciprocal architecture (blue line) in panels A and B broadly captures the wildtype NPAO1 behavior reported in Rattray et al. (2022). Compared to this baseline, the figure predicts that removing the influence of *rhl* on *las* (hierarchical architecture, dark green and red lines) results in a delayed response to increasing density. Removing all *las/ rhl* interactions (independent architecture, light green and orange lines) significantly attenuates and delays the response to density. The reciprocal architecture is the most sensitive to changes in population density as its *lasB* expression fold-change increases the most for a given density value.

Figures 10C and 10D examine the impact of different architectures on the bacteria’s response to the physical environment. In this case we show expression level as a function of processes affecting mass transfer such as advective flow or diffusive loss of signal. Here, the reciprocal architecture is the least sensitive to changes as its *lasB* expression has the smallest decline for a given mass transfer rate.

## Discussion

In this study we consider different architectures for multi-signal quorum sensing systems (Figure 1) and show that the conventional *las-rhl* hierarchical view of QS in *P. aeruginosa* (Figure 2) is incomplete. Specifically, we find that both the *las* and *rhl* systems regulate each other. Figure 3 corroborates the influence of *las* on *rhl*, but, contrary to the hierarchical view, we also show in Figure 4 that the *rhl* signal C_4_-HSL can influence the *las* synthase *lasI*. This effect is significant, as C_4_-HSL alone induces more than a six-fold increase in *lasI* expression compared to basal levels. We confirm these results when both signals are present simultaneously (Figure 5), and further show that both *las* and *rhl* synthase genes require both signals for maximal expression. By fitting a mathematical model, we demonstrate that simple additive effects are insufficient to explain our data (Figure 6). Closing the gap apparent in Figure 6 requires that the signals interact cooperatively to augment their additive effects. By modeling both the reciprocal relationship and the effect of cooperativity, we provide a quantitative model for QS in a model system, and conclude that the *las*-*rhl* relationship forms a biased reciprocal network (Figure 7). We then model the effect of this architecture on a representative QS-controlled gene (*lasB*) and compare the results with other architectures. By extending existing dynamical system models we estimate the different steady state signal concentrations (Figure 8) and use those estimates to predict *lasB* expression. We find that the reciprocal architecture is more sensitive to population density and more robust in the presence of environmental interference (Figure 10).

By focusing on signal concentration as the factor determining behavior, our approach accommodates multiple possible molecular mechanisms. It does mean, however, that we cannot easily distinguish between them. For example, C_4_-HSL could be causing an increase in *lasI* expression by enabling the formation of LasR dimers, albeit less efficiently than 3-oxo-C_12_-HSL. Alternatively, it could be the case that the RhlR/C_4_-HSL complex serves as an activating transcription factor for rhlI. Additional experiments would be required to distinguish between these two cases.

It should be noted that our use of *ΔlasIΔrhlI* cells might cause differences between our observations and wild type responses. In particular, we make two assumptions about the mutant strain. First, we assume that the only effect of the *lasI* and *rhlI* deletions is an inability to successfully produce LasI and RhlI proteins. Secondly, we assume that the only relevant phenotypic function of those proteins is the synthesis of the corresponding signal molecules. Although we cannot rule out pleiotropic effects from the strain construction or lack of synthase proteins, we do not expect that any such effects would alter our conclusions.

Our first key result demonstrates that the *las* and *rhl* systems form a reciprocal architecture, extending existing research into the relationship between those systems. Many researchers, including Pesci et al. (1997), de Kievit et al. (2002), and Medina et al. (2003), have shown that the *las* system is essential for maximal expression of genes in the *rhl* system. Our data substantiates those results, but we also show the converse: the *rhl* system, in particular its signal C_4_-HSL, is essential for maximum expression of a gene in the *las* system. We further extend prior results by considering the combination of both signals and by quantifying the relationship between the systems. Most previous attempts at this quantification have assumed a hierarchical architecture. For example, de Kievit et al. (2002) demonstrate that LasR/3-oxo-C_12_-HSL alone influences *rhlI* expression more than RhlR/ C_4_-HSL alone, a result consistent with our data. Their analysis, however, is limited to measuring the response of the *rhl* system. Analyses of the effect of C_4_-HSL on the *las* system are much less common, though Wargo and Hogan (2007) do report that a *rhlI* mutant produced the same 3-oxo-C_12_-HSL concentrations as wild type. Those experiments were conducted in *Escherichia coli*, however, and the authors acknowledge that *E. coli* may include its own regulators that mimic the behavior of C_4_-HSL. As with other published reports, the focus is on single signals in isolation, which necessarily neglects the effect of both signals combined.

In shifting from the molecular to the population level, we adopt the single signal QS models of Dockery and Keener (2001) and Brown (2013). This approach has produced both theoretical conjectures (Pai and You 2009) and experimental interpretations (Fekete 2010). Our analysis extends the models to account for multiple QS signals and interactions between them. In this way we can study not only isolated quorum sensing systems, but also networks of interrelated systems. We can both characterize the architectures of those systems and quantify the intra-system and inter-system effects.

Consideration of multi-system architects can lead to insights in bacterial population-level behavior. Combinatorial quorum sensing (Cornforth et al. 2014), for example, postulates that having two different QS systems allows populations to discriminate between social (high density) and physical (high containment) environments. In particular, if the two signals have different decay rates, then their concentrations will respond differently to variation in population density, while other environmental conditions such as mass transfer will effect each signal equivalently. Downstream genes whose expression depends on a combination of both signals can then differentiate between changes in the two environmental dimensions. The authors measure a significant difference in the decay rates of C_4_-HSL and 3-oxo-C_12_-HSL, and they identify sets of downstream genes whose activation are controlled by different combinations of multi-signal inputs (e.g. Boolean and, or, logic). Their analysis, however, only considers an architecture in which the *las* and *rhl* QS systems are independent.

The model developed from equation 3 allows us to return to combinatorial quorum sensing and evaluate it using defined and parameterized QS architectures. Following Cornforth et al. we plot isoclines of the C_4_-HSL and 3-oxo-C_12_-HSL signal concentrations as functions of the same environmental dimensions, population density, *N*, and mass transfer rate, *m*. Figure 11 shows the results for the independent architecture Cornforth et al. assume (Figure 11A) and for the reciprocal architecture we have established (Figure 11B). The parameter values from Tables 2 and 3 were again used to construct both plots. As expected from Figure 7, the reciprocal architecture has a larger region in which either QS system is active (Boolean or); however, the fraction of that region in which *both* QS systems are active (Boolean and) is lower with a reciprocal architecture. In this example a reciprocal architecture provides greater discrimination between the two environmental dimensions.

**Figure 11.**
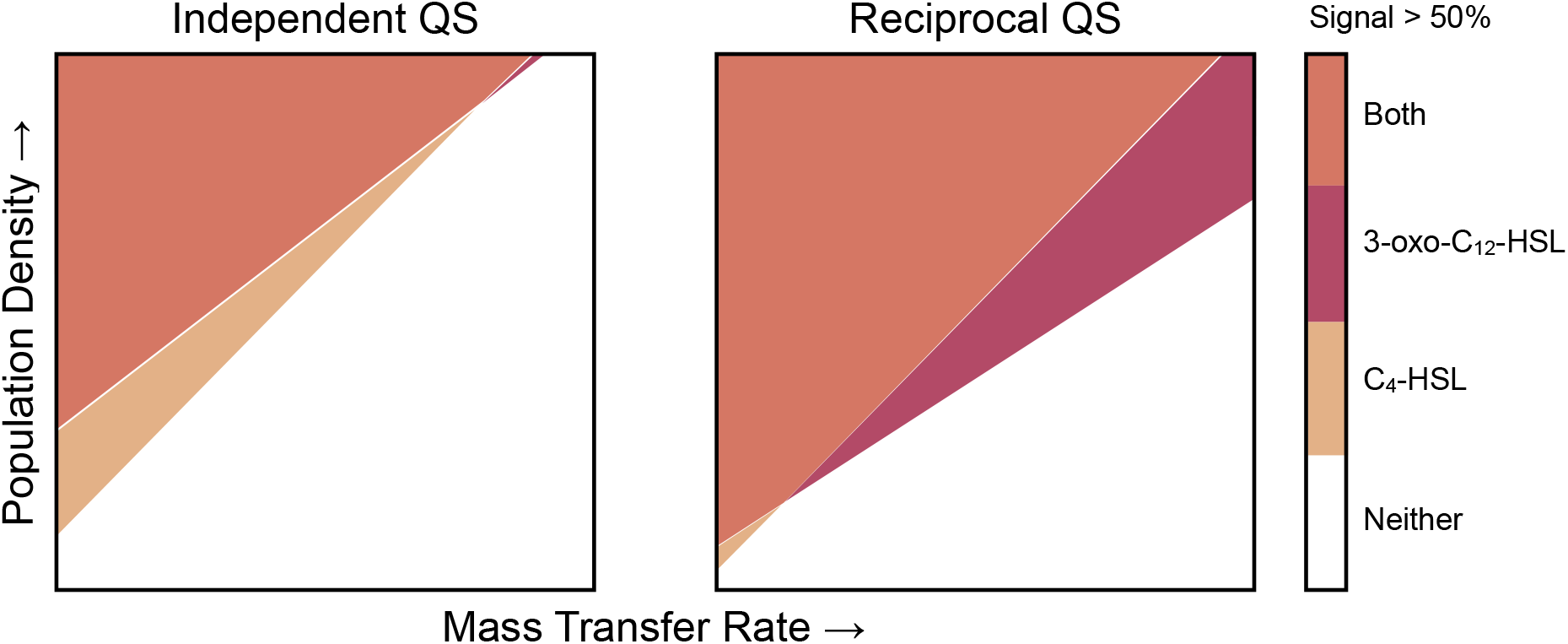
A reciprocal QS architecture can increase the discriminatory power of combinatorial quorum sensing. Each panel shows the environmental and social regions in which Boolean combinations of individual QS systems are active. A QS system is considered active if its associated signal concentration exceeds 50% of its maximum value. In both panels density ranges from approximately 0.5 OD to 1.0 OD, while normalized mass transfer rate ranges from 0 to the decay rate of C_4_-HSL. With the independent architecture, the Boolean and condition comprises 80% of all QS activation; with the reciprocal architecture, that percentage is reduced to 75%.

By choosing the *las* and *rhl* quorum sensing systems for this study, we focus on a relatively straightforward QS architecture. It consists of only two systems, and the two systems reinforce each other. Our approach is not limited to this case, however. Equations 2 and 3 can accommodate more than two systems and account for systems that repress expression as well as enhance it. Both enhancements are relevant for our model organism, as *Pseudomonas aeruginosa* includes additional QS systems beyond *las* and *rhl;* there is also the *pqs* system (Pesci et al. 1999). And *P. aeruginosa* is not unique in having multiple QS systems (Papenfort and Bassler 2016). Two frequently studied species (*P. aeruginosa* and *V. harveyi*) each have at least three parallel systems (Ng and Bassler 2009), and some may have as many as eight (Brachmann et al. 2013). Furthermore, different systems do not always reinforce each other. In some cases one system can repress another (McGrath et al. 2004). We hope to address both considerations in future work, as even a simple model shows potential for providing important insights. Consider the case where one QS system, on activation, represses expression in another. That repression can limit expression of the other system, or it can stop expression entirely. Those possibilities, illustrated in Figure 12, may have vastly different effects on the population-level behavior in response to changes in stationary phase density.

**Figure 12.**
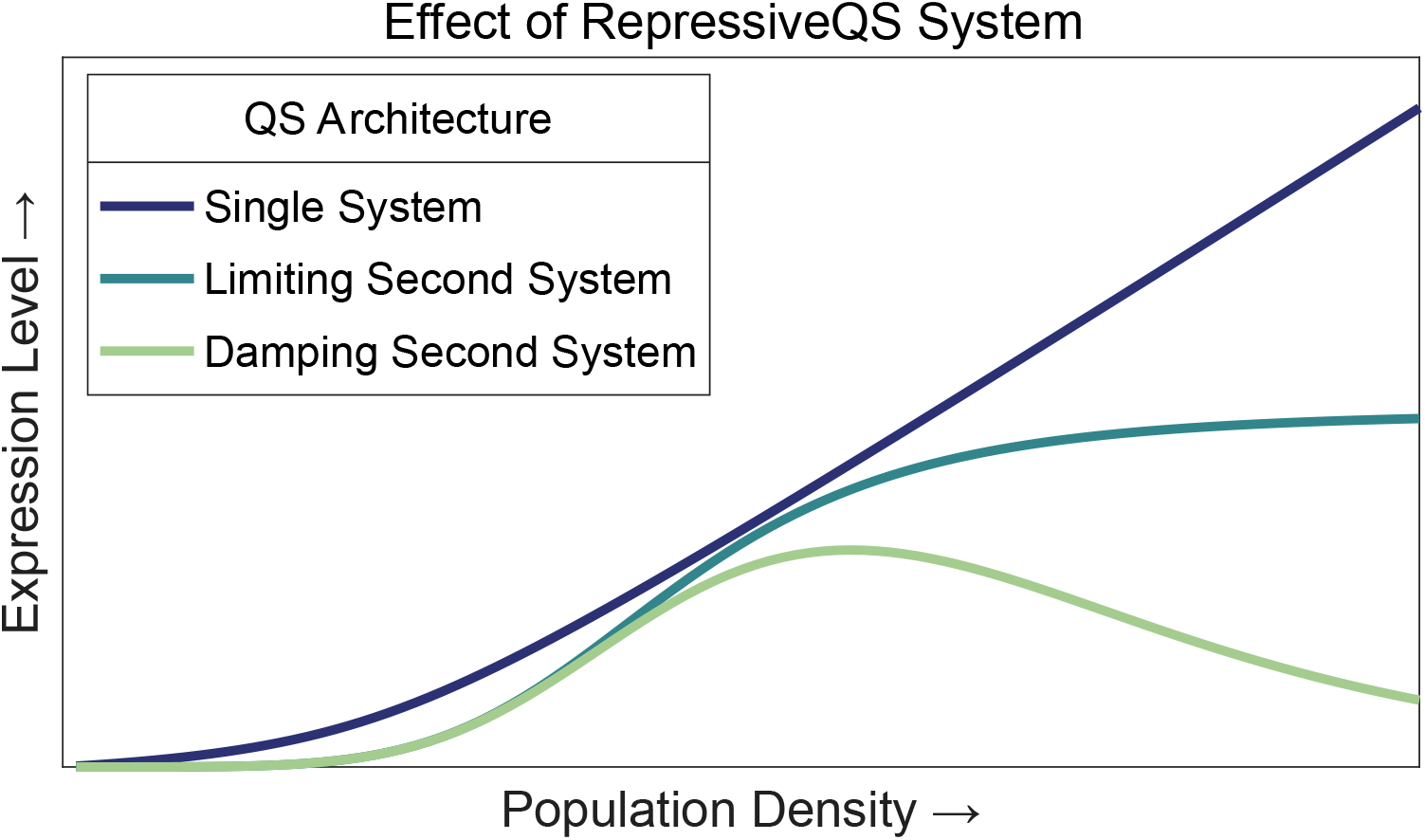
Repressive quorum sensing systems can have various effects on the overall response. The plot shows the overall response (e.g. expression level of a downstream gene) to population density for two types of repressive systems. It also shows the response of an unconstrained single system for comparison.

Finally, although we are able to model QS architectures at the cellular and population level, it is not clear how traditional gene regulatory networks can achieve the responses we observe. Long et al. (2009) suggest that multiple activating transcription factors combine additively, but that can only be true if the effects of each are independent. In contrast, Kaplan et al. (2008) claim that multiple inputs controlling gene expression usually combine multiplicatively. This relationship holds when the binding of one factor to the promoter depends on the presence of the second at that promoter. As Figure 6 makes clear, neither approach can adequately explain our data.

Sauer et al. (1995) make related observations for a protein complex in *Drosophila melanogaster*; both of the developmental regulators BCD and HB alone induce a 6-fold increase by themselves but combine to induce a greater than 65-fold increase. Their results offer a tantalizing possibility that further investigations into the mechanisms of *P. aeruginosa* quorum sensing interactions can provide insights into more general gene regulatory networks.

## Methods

### Data Collection

We used three strains for the experimental observations: l*asB:luxCDABE* genomic reporter fusion in NPAO1*ΔlasIΔrhlI, lasI:luxCDABE* genomic reporter fusion in NPAO1*ΔlasIΔrhlI*, and *rhlI:luxCDABE* genomic reporter fusion in NPAO1*ΔlasIΔrhlI*. We streaked out all strains in Luria-Bertani (LB) agar at 37°C for 24 hours and then subcultured a single colony in 10 mL LB, incubated at 37°C under shaking conditions (180 rpm) for 24 hours.

We prepared 3-oxo-C_12_-HSL and C_4_-HSL in methanol at 7 different concentrations: 0.1, 0.5, 1, 2, 3, 4 and 5 μM, each diluted from 100 mM stock. We centrifuged all cultures and washed each three times using PBS. We then re-suspended in LB and diluted to an OD (600) of 0.05. We then transferred 200 μl of each culture to a black 96-well plate with a clear bottom and inoculated with signals at the indicated concentrations. We repeated each experiment to generate five replicates. Methanol with no signal was used as a control. The plates were incubated in BioSpa at 37°c for 18 h. Measurements of OD (600) and RLU (Relative Luminescence Units) were collected every hour.

### Mathematical Models

Parameters for the kinetic models of equations 1 and 2 were estimated by minimizing sum of squared errors. Observations were limited to time ranges with peak expression values. (See Supplementary Information for detailed time course analysis.) Figure SI.4 compares the predictions from equation 1 model with experimental observations. Figure SI.5 compares the predicted expression levels from equation 2 model and observed values.

More detailed comparisons are available in Supplementary Information.

## References

Bolouri, Hamid. Computational Modeling of Gene Regulatory Networks: A Primer. Imperial College Press, 2008.

Brachmann, AO, S Brameyer, D Kresovic, I Hitkova, Y Kopp, C Manske, K Schubert, HB Bode, and R Heermann. “Pyrones as Bacterial Signaling Molecules.” Nat Chem Biol 9, no. 9 (2013): 573–78.

Brown, D. “Linking Molecular and Population Processes in Mathematical Models of Quorum Sensing.” Bull Math Biol 75, no. 10 (2013): 1813–39.

Buchler, NE, U Gerland, and T Hwa. “On Schemes of Combinatorial Transcription Logic.” Proc Natl Acad Sci U S A 100, no. 9 (2003): 5136–41.

Casilag, F, A Lorenz, J Krueger, F Klawonn, S Weiss, and S Haüssler. “The Lasb Elastase of Pseudomonas Aeruginosa Acts in Concert With Alkaline Protease Apra to Prevent Flagellin-Mediated Immune Recognition.” Infect Immun 84, no. 1 (2016): 162–71.

Cigana, Cristina, Jérôme Castandet, Nicolas Sprynski, Medede Melessike, Lilha Beyria, Serena Ranucci, Beatriz Alcalá-Franco, Alice Rossi, Alessandra Bragonzi, and Magdalena Zalacain. “Pseudomonas Aeruginosa Elastase Contributes to the Establishment of Chronic Lung Colonization and Modulates the Immune Response in a Murine Model.” Frontiers in Microbiology 11 (2021): 620819.

Cornforth, DM, R Popat, L McNally, J Gurney, TC Scott-Phillips, A Ivens, SP Diggle, and SP Brown. “Combinatorial Quorum Sensing Allows Bacteria to Resolve Their Social and Physical Environment.” Proc Natl Acad Sci U S A 111, no. 11 (2014): 4280–84.

de Kievit, TR, Y Kakai, JK Register, EC Pesci, and BH Iglewski. “Role of the Pseudomonas Aeruginosa Las and Rhl Quorum-Sensing Systems in Rhli Regulation.” FEMS Microbiol Lett 212, no. 1 (2002): 101–6.

Diggle, SP, AS GrifYin, GS Campbell, and SA West. “Cooperation and ConYlict in Quorum-Sensing Bacterial Populations.” Nature 450, no. 7168 (2007): 411–14.

Dockery, JD, and JP Keener. “A Mathematical Model for Quorum Sensing in Pseudomonas Aeruginosa.” Bull Math Biol 63, no. 1 (2001): 95–116.

Fekete, A, C Kuttler, M Rothballer, BA Hense, D Fischer, K Buddrus-Schiemann, M Lucio, J Müller, P Schmitt-Kopplin, and A Hartmann. “Dynamic Regulation of N-Acyl-homoserine Lactone Production and Degradation in Pseudomonas Putida Isof.” FEMS Microbiol Ecol 72, no. 1 (2010): 22–34.

James, S, P Nilsson, G James, S Kjelleberg, and T Fagerström. “Luminescence Control in the Marine Bacterium Vibrio Fischeri: An Analysis of the Dynamics of Lux Regulation.” J Mol Biol 296, no. 4 (2000): 1127–37.

Jayakumar, P, ART Figueiredo, and R Kümmerli. “Evolution of Quorum Sensing in Pseudomonas Aeruginosa Can Occur Via Loss of Function and Regulon Modulation.” mSystems 7, no. 5 (2022): e0035422.

Kaplan, S, A Bren, A Zaslaver, E Dekel, and U Alon. “Diverse Two-Dimensional Input Functions Control Bacterial Sugar Genes.” Mol Cell 29, no. 6 (2008): 786–92.

LatiYi, A, M Foglino, K Tanaka, P Williams, and A Lazdunski. “A Hierarchical Quorum-Sensing Cascade in Pseudomonas Aeruginosa Links the Transcriptional Activators Lasr and Rhir (Vsmr) to Expression of the Stationary-Phase Sigma Factor Rpos.” Mol Microbiol 21, no. 6 (1996): 1137–46.

Long, T, KC Tu, Y Wang, P Mehta, NP Ong, BL Bassler, and NS Wingreen. “Quantifying the Integration of Quorum-Sensing Signals With Single-Cell Resolution.” PLoS Biol 7, no. 3 (2009): e68.

McGrath, S, DS Wade, and EC Pesci. “Dueling Quorum Sensing Systems in Pseudomonas Aeruginosa Control the Production of the Pseudomonas Quinolone Signal (Pqs).” FEMS Microbiol Lett 230, no. 1 (2004): 27–34.

Medina, G, K Juárez, R Díaz, and G Soberón-Chávez. “Transcriptional Regulation of Pseudomonas Aeruginosa Rhlr, Encoding a Quorum-Sensing Regulatory Protein.” Microbiology (Reading) 149, no. Pt 11 (2003): 3073–81.

Ng, WL, and BL Bassler. “Bacterial Quorum-Sensing Network Architectures.” Annu Rev Genet 43 (2009): 197–222.

Nouwens, AS, SA Beatson, CB Whitchurch, BJ Walsh, HP Schweizer, JS Mattick, and SJ Cordwell. “Proteome Analysis of Extracellular Proteins Regulated By the Las and Rhl Quorum Sensing Systems in Pseudomonas Aeruginosa Pao1.” Microbiology (Reading) 149, no. Pt 5 (2003): 1311–22.

Pai, A, and L You. “Optimal Tuning of Bacterial Sensing Potential.” Mol Syst Biol 5 (2009): 286.

Papenfort, K, and BL Bassler. “Quorum Sensing Signal-Response Systems in Gram-Negative Bacteria.” Nat Rev Microbiol 14, no. 9 (2016): 576–88.

Pearson, JP, EC Pesci, and BH Iglewski. “Roles of Pseudomonas Aeruginosa Las and Rhl Quorum-Sensing Systems in Control of Elastase and Rhamnolipid Biosynthesis Genes.” J Bacteriol 179, no. 18 (1997): 5756–67.

Pesci, EC, JB Milbank, JP Pearson, S McKnight, AS Kende, EP Greenberg, and BH Iglewski. “Quinolone Signaling in the Cell-to-cell Communication System of Pseudomonas Aeruginosa.” Proc Natl Acad Sci U S A 96, no. 20 (1999): 11229–34.

Pesci, EC, JP Pearson, PC Seed, and BH Iglewski. “Regulation of Las and Rhl Quorum Sensing in Pseudomonas Aeruginosa.” J Bacteriol 179, no. 10 (1997): 3127–32.

Rattray, JB, SA Thomas, Y Wang, E Molotkova, J Gurney, JJ Varga, and SP Brown. “Bacterial Quorum Sensing Allows Graded and Bimodal Cellular Responses to Variations in Population Density.” mBio 13, no. 3 (2022): e0074522.

Santillán, Moises. “On the Use of the Hill Functions in Mathematical Models of Gene Regulatory Networks.” Mathematical Modelling of Natural Phenomena 3, no. 2 (2008): 85–97.

Sauer, F, SK Hansen, and R Tjian. “Multiple TaYiis Directing Synergistic Activation of Transcription.” Science 270, no. 5243 (1995): 1783–88.

Schuster, M, and EP Greenberg. “Early Activation of Quorum Sensing in Pseudomonas Aeruginosa Reveals the Architecture of a Complex Regulon.” BMC Genomics 8 (2007): 287.

Sexton, DJ, and M Schuster. “Nutrient Limitation Determines the Fitness of Cheaters in Bacterial Siderophore Cooperation.” Nat Commun 8, no. 1 (2017): 230.

Stearns, SC. “The Evolutionary SigniYicance of Phenotypic Plasticity.” Bioscience (1989):

Ward, JP, JR King, AJ Koerber, P Williams, JM Croft, and RE Sockett. “Mathematical Modelling of Quorum Sensing in Bacteria.” IMA J Math Appl Med Biol 18, no. 3 (2001): 263–92.

Wargo, MJ, and DA Hogan. “Examination of Pseudomonas Aeruginosa Lasi Regulation and 3-Oxo-c12-homoserine Lactone Production Using a Heterologous Escherichia Coli System.” FEMS Microbiol Lett 273, no. 1 (2007): 38–44.

Wellington, S, and EP Greenberg. “Quorum Sensing Signal Selectivity and the Potential for Interspecies Cross Talk.” mBio 10, no. 2 (2019): e00146–19.

